# Drought Resistance of 10 Ground Cover Seedling Species during Roof Greening

**DOI:** 10.1101/711051

**Authors:** Pengqian Zhang, Jiade Bai, Yanju Liu, Yuping Meng, Zheng Yang, Tian Liu

## Abstract

10 species’ drought resistance cases have been studied, including *Paeonia lactiflora*, *Hemerocallis dumortieri*, *Physostegia virginiana*, *Iris lacteal*, *Hylotelephium erythrostictum*, *Sedum lineare*, *Iris germanica*, *Cosmos bipinnata*, *Hosta plantaginea* and *Dianthus barbatus*. By researching these drought resistance cases, a suggestion can be given for roof greening. This research sets 3 drought stress scenarios by controlling the soil relative water content (RWC), including moderately drought stress (40%±2% < RWC < 45%±2%), strong drought stress (RWC < 30%±2%) and control group (RWC > 75%±2%). After the seedlings survived the drought stress, the damaging rate of permeability (DRP), total chlorophylls concentrations (Chl), superoxide dismutase (SOD), peroxidase (POD) and ascorbate peroxidase (AsAPOD) of seedlings will be measured. Finally, a subordinate function method was applied to assess these species’ drought resistance. *Cosmos bipinnata* and *Physostegia virginiana* was dead after having suffered with moderately drought stress and strong drought stress, respectively. Although other species survived, the individual variation was huge especially for physiological and biochemical index. *Hemerocallis dumortieri*, *Iris lactea* and *Hosta plantaginea*’s DRP had little change when they lived in the normal water condition and suffered with drought stress. Most of the species (except *Paeonia lactiflora* and *Sedum lineare*) showed a lower SOD activity during moderately drought stress compared with the sufficient soil water condition and strong drought stress condition. The changes of plants’ POD activity and AsAPOD activity are very similar: when drought stress enhanced, the activity of protect enzyme reduced. According to the subordinate function method, the order of plants’ resistance to the drought is as follow: *Hosta plantaginea* > *Sedum lineare* > *Iris germanica* > *Hemerocallis dumortieri* > *Iris lactea* >*Hylotelephium erythrostictum* > *Dianthus barbatus* > *Paeonia lactiflora* > *Physostegia virginiana* > *Cosmos bipinnata*.

## 1 Introduction

Roof greening, regarded as “Fifth surface greening”, is one of the fundamental measures for sponge city, a significant national policy to improve relationships between city development and nature protection, in order to keep hydro-ecology balance (Wang and Zhang, 2015; Yu *et al*., 2015). As an important supplement of urban landscaping, roof greening is good for mitigating Urban Heat Island Effect (UHI) (Ondimu and Murase, 2007), improving air quality (Baik *et al*., 2012) and enriching biodiversity of city (Köhler and Clements, 2013), hence the reason to extend such landscaping style to all of China. Green roof can be roughly categorized into two types: support diverse plants (shrubs and trees, grass and flowers), namely intensive green roof (IGR) and simple herbaceous plant species, namely extensive green roof (EGR) (Peng and Jim, 2015). The plant has many challenges to grow on the roof. Take Beijing as an example, plants applied to the roof green are suffered with restricted rainfall in the winter, spring and autumn; the evaporation always increased by summer high temperature. Drought is considered one of the most common environmental stresses presently affecting plant growth (Nahar *et al*., 2015; Romain *et al*., 2006). When plants are suffering drought stress, the reactive oxygen species (ROS) would produce in the plant (Sharma and Dubey, 2005). Different ROS, including singlet oxygen (^1^O_2_), superoxide radical (O_2_ ^−^), hydroxyl free radical (•OH), and hydrogen peroxide (H_2_O_2_) (Smirnoff, 1993) will reduce the productivity in crop and depress viability in the plant, as they would cause oxidative damage to proteins, DNA, and lipids (Apel and Hirt, 2004). Accordingly, drought stress will disturb leaf membrane permeability (MP) (Bai *et al.*, 2006), the total chlorophyll concentrations (Chl) (Rulcová and Pospíšilová, 2001), superoxide dismutase (SOD) (Bedard and Krause, 2007), peroxidase (POD) (Bahari *et al*., 2015), and ascorbate peroxidase (AsAPOD) (Ghozlene *et al*., 2014). Actually, these indicators mentioned above that measure the degree of plants bearing drought stress include MP, Chl, SOD, POD and AsAPOD usually are analyzed as a whole. That is because these indexes have strong relations with each other, for instance SOD, POD and AsAPOD as anti-oxidative enzymes (Chang *et al*., 2006) can eliminate ROS to protect the cells from being damaged.

According to the “Beijing local standards, roof greening specification (DB11/T 281-2005)” (Beijing Municipal Administration of Quality and Technology Supervision), there are more than 20 ground cover seedlings included *Paeonia lactiflora* Pall., etc., that recommended to applying during the roof greening. However, these plants’ drought resistance has not sequenced. To figure out the plant with strong resistance to the drought environment can improve the surviving rate of these plants and save maintenance cost during roof greening.

## 2 Study Area Overview

Milu Park is situated 2 km to the south of the South-5-Ring Road in Beijing, and surrounded by Nan-Haizi Suburb Park. The drought stress test was held in a rain protection shed, located in the core-protection area of David’s Deer in Milu Park (39.78N, 116.47E). The daily average temperature was about 18.7 ℃, the daily average humidity was about 55.2%, the daily average illumination intensity (at 12:00) was about 2000 Lx, during the test period. The test was held at the end of April to the end of May, 2015.

## 3 Materials and methods

### 3.1 Plant seedlings

10 first-year seedlings include *Paeonia lactiflora* Pall., *Hemerocallis dumortieri* Morr., *Meehania urticifolia* (Miq.) Makino, *Iris lactea* Pall. var. *chinensis* (Fisch.) Koidz., *Hylotelephium erythrostictum* (Miq.) H. Ohba, *Sedum lineare* Thunb., *Iris germanica*, *Cosmos bipinnata* Cav., *Hosta plantaginea* (Lam.) Aschers. and *Dianthus barbatus* L. are chosen as materials to be tested under different drought stress degrees. These plants have different propagation modes and other characteristics, the details see Table 1. These seedlings were provided by Yu-quanying flowers market, a big and famous market in Beijing.

**Table 1.**
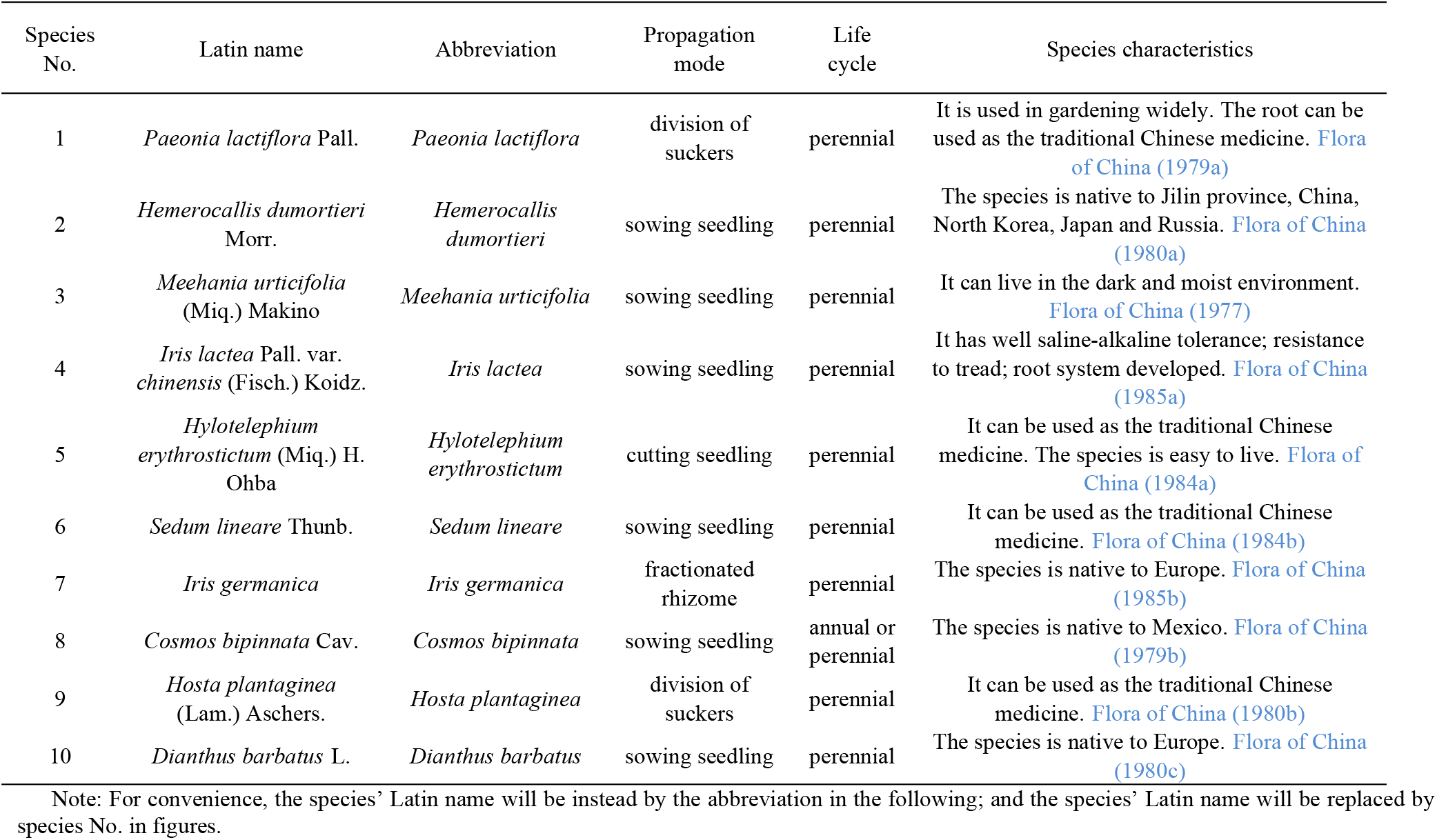
Plant species list and basic features at the three sampling sites

### 3.2 Field soil collection

The plant seedlings need replant into the special flowerpot, that was plastic made, with 20 cm high and 13 cm diameters, and there are 3 little holes at the bottom of the pot for pervious to water for the drought experiment. The field soil in Tao-huadao, a small wetland already dried up entirely in Milu Park was used as soil for transplanting. The soil is belonging to the medium salty soil, with mean value of pH are 7.89, mean content rapidly available nitrogen are 24.73 mg⋅kg^−1^, mean content rapid available phosphorus are 18.87 mg⋅kg^−1^, mean content rapidly available potassium are 322.18 mg⋅kg^−1^ (Zhu *et al.*, 2016). The field soil was pulverized to powders by rolling stick after they dried out naturally.

### 3.3 Plants transplanting

The program of transplantation is as follows: firstly, about 400 g soil powder was put in the pot. Secondly, tear off the one-off plastic wrap surrounded on the seedlings’ roots. Then pick up the seedlings very carefully, and put them in the pot. Place the roots as close to the center of the pot as possible. Thirdly, fill other about 300 g soil powder into the pot and bury the root. Those soil powders need compressed tightly with fingers. At last, water seedlings once every 10 min and repeat 3 times, for keeping there are enough water support the plant rooting. The success of transplanting a plant can be judged by observing whether it grows new leaves, and whether the stem is fresh. After a week of replanting, the seedlings’ surviving rate achieved 99%.

### 3.4 Drought stress design

Three drought stress degrees have been set, including moderately drought stress (Treatment 1 is short for *T1*, the water content in soil varies from 40%±2% ~ 45%±2%), strong drought stress (Treatment 2 is short for *T2*, the water content in soil are less than 30%±2%) and sufficient soil water condition (Control group is short for *Cg*, the water content in soil are higher than 75%±2%) (Bai *et al*., 2015; Bu *et al*., 2010). For every drought stress degrees there were 30 seedlings with 3 replicates for each plant species have been tested.

For Cg, water plant seedlings every 4 days. The drought stress was dependent on natural evaporation. During the drought stress period, the W.E.T Sensor type WET-2^TM^ made by Delta-T Devices Limited Company was used to measure the water content. Once the relative water content of soil meets the requirements of experiment, keep the plant seedlings lived in such environment about 2 days to make sure the physiology and biochemistry have been changed.

### 3.5 Leaves sampling

The details of sampling method in VDI-Guideline 3975 Part 11 (2007) are referenced. Leaves of 10 plant species were collected during April 27^th^ to May 9^th^, 2015. Triplicate samples were collected for each plant species of each treatment. At least 15 g of varies numbers of leaves has been collected for each sample in order to have enough quantity for analysis. Healthy-looking leaves were chosen as possible. Leaves were placed into sealed plastic bags and kept in a portable ice-box at 0 ~ 4 ℃ before being transferred to the lab for further physiological and chemical analysis.

### 3.6 MP, Chl, SOD, POD and AsAPOD determination

Membrane Permeability of leaves was determined as 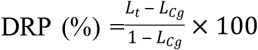. Where *L*_*t*_ is the relative electrical conductance of drought stress treatments; *L*_Cg_ is the relative electrical conductance of control group. The relative electrical conductance followed 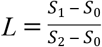 Where *S*_1_ is the original conductivity value of the deionized water with fractured fresh leaves; *S*_2_ is the conductivity value of the boiled deionized water with fractured leaves; *S*_0_ is the conductivity value of deionized water (Zhang *et al*., 2011). Leaves membrane permeability was determined using a model Thermo^TM^ Scientific Orion 3-star inductivity measurer. Before the test, all samples were flushed by de-ionized water for 3 times and wipe off water on the leaves’ surface.

The total chlorophyll content estimation was carried out by the method of Arnon (1949). The method details were described in a published paper (Zhang *et al.*, 2016).

Treating the crude enzyme extracted from leaves is the impurity to measure the SOD, POD and AsAPOD. About 0.5 g fresh leaves with a small number of CaCO_3_ and high purity quartz sand were crashed into powder under freezing environment, with 5 mL phosphate buffer (0.05 mol⋅L^−1^) in a mortar. The paste were breathed into a 10 mL centrifuge tube, and then diluted with de-ionized water to 10 mL. By centrifuging at high speed (F=1 3000 g) for 20 min at 0 ℃ to 4 ℃ (Zhang *et al*., 2011).

SOD and POD reaction system was described by Zhang *et al* (2011), AsAPOD reaction systems was described by Tang *et al* (2012) as follows:

Based on the reaction system (Table 2-SOD) mentioned above, every sample was put into test tube and shines them under 4000 Lx illumination intensity about 20-30 min at the room temperature except the *Illumination Check*. Once color of solution transition started, the reaction should be stop immediately. The final solution absorbance value was determined with METASH™ UV-6100A Spectrophotometer at 560 nm wavelength.

**Table 2.**
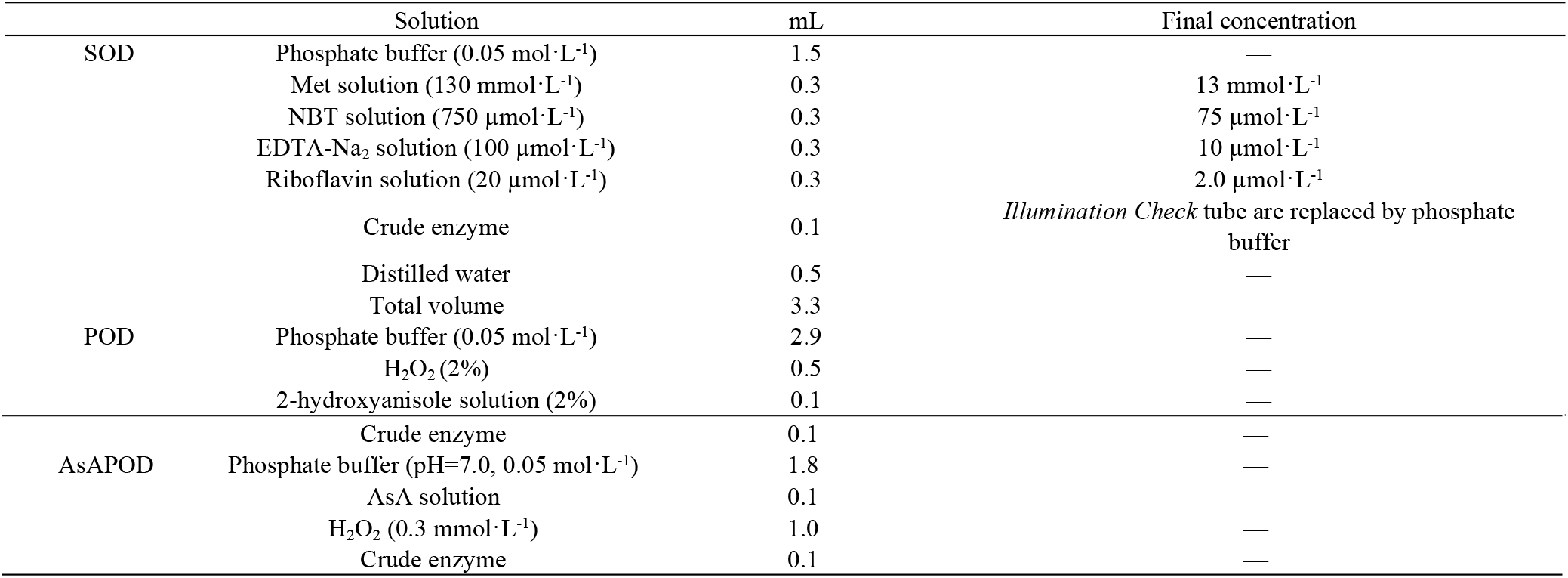
The solution of SOD, POD and AsAPOD reaction system

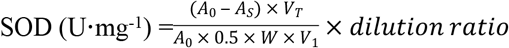

Where *A*_0_ is the absorbance of *Illumination Check*, when all the tubes shined with the light, the check tube is wrapped in the tinsel avoid be shined; *A*_s_ is the is the absorbance of samples; *V*_T_ (mL) is the total volume of samples; *V*_1_ (mL) is the volume of participation in the reaction and *W* (g) is the weight of the fresh leaves.

All the solution were put into the test tube, which followed by the POD reaction system (Table 2-POD). Then record the solution’s light absorption value of each tube (the wave length was regulated at 470 nm). Read every 1 minute, and each solution was recorded 5 times in 5 minutes.

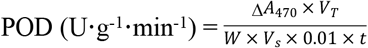

Where Δ*A*_470_ is the change of absorbance during the reaction period; *W* (g) is the weight of the sample; *t* (min) is the reaction time; *V*s (mL) is the volume of participation in the reaction and the *V*_T_ (mL) is the total volume of samples.

According to the AsAPOD reaction system (Table 2-AsAPOD), the mixture was put into test tubes, and then records the light absorption values at 290 nm at every minute. And such the step repeats 5 times. The formula to account the AsAPOD as follows:

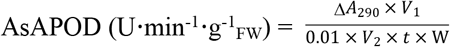

Where Δ*A*_290_ is the change of absorbance during the 5 min; *V*_1_ (mL) is the volume of crude enzyme; *V*_2_ (mL) is the volume of crude enzyme to take part in the reaction (0.1 mL in this test); *t* (min) is the reaction time (5 min in this test) and the *W* (g) is the weight of the fresh leaves. FW is the short for leaves fresh weight.

### 3.7 Data analysis

SPSS 17.0 and Excel 2010 for Windows were applied to calculate the mean, S.D., etc. A multiple comparison of the means by least significant difference (Tukey HSD) test was performed to the 5 indexes (DRP, Chl, SOD, POD and AsAPOD) under the 2 drought stress levels and control group. And ANOVA indicated significant levels as (*P*<0.01; 0.05).

A drought resistance assessment method (Wu *et al*., 2013; Chen *et al*., 2002) for the plant species based on the subordinate function value belong to the fuzzy mathematics is applied in the research. By the method, the DRP, Chl, SOD, POD and AsAPOD’s results can be synthesized as a single value of each species. The subordinate function value formula is as follows:

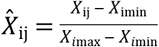

Where the lowercase “i” and “j” is represents plant species and determination index respectively. Therefore, the “*X*_ij_” is the mean value of “j” index of the “i” species. “*X*_imax_” and “*X*_imin_” is represents the maximums and minimums of the “j” index of the “i” species. 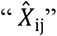 is the subordinate function value. 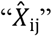 represents the drought resistance of plant seedlings. Then the average of subordinate function value is applied to estimate the adaptive capacity of plants that bear the drought stress. The average formula is as follows (“*n*” represents the amount of indexes and 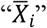 represents the average):

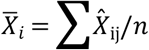

## 4 Results and discussions

### 4.1 Soil moisture content

The water content of soil sees Table 3. Although the water content of soil has standard deviation, the 3 drought stress levels are distinguished, well agreeable to the designed levels.

**Table 3.** The moisture content of soil

| Plant species | Cg (%) | T1 (%) | T2 (%) |
| --- | --- | --- | --- |
| *Paeonia lactiflora* | 80%±2% | 45%±4% | 20%±2% |
| *Hemerocallis dumortieri* | 82%±4% | 41%±2% | 17%±4% |
| *Physostegia virginiana* | 79%±5% | 41%±3% | 17%±2% |
| *Iris lactea* | 85%±1% | 42%±3% | 21%±3% |
| *Hylotelephium erythrostictum* | 83%±2% | 40%±3% | 21%±2% |
| *Sedum lineare* | 78%±1% | 43%±1% | 24%±3% |
| *Iris germanica* | 85%±2% | 44%±2% | 22%±2% |
| *Cosmos bipinnata* | 77%±2% | 40%±3% | 20%±4% |
| *Hosta plantaginea* | 84%±3% | 42%±2% | 23%±3% |
| *Dianthus barbatus* | 82%±4% | 40%±1% | 24%±1% |

Note: Values represent mean±SD ( $n=30$ ). Cg is short for control group (with normal watering condition); $T_1$ represents the “moderately drought stress”; $T_2$ represents the “strong drought stress”. Greek alphabet “I, II and III” represents the first, second and third drought stress tests.

### 4.2 Membrane permeability

The values of MP represent how much the electrolyte seeping from the cell of plant. According to Sanchez-Rodriguez *et al*. (2010), the maintenance of sufficient water in plant tissue protects plant from dehydration and carboxylation, and their other enzymes from inactivation. The low-temperature environment (Shen, 2005), drought stress environment (Dai *et al*., 2006), salt stress environment (Xu *et al*., 2002; Du *et al*., 2012) and heavy metal stress (Vos *et al*., 1989) has been proven to achieve a higher level of damaging rate of permeability.. In other words, the higher value of MP is, the more the cell damage is. With the same water condition, the lower MP value implies the stronger adaptation of plant species to the environment (Wang *et al*., 2014).

MP values between different plant species under control treatment (*Cg*) show the plant itself living status. The MP mean value from high to low as *Cosmos bipinnata*, *Dianthus barbatus*, *Physostegia virginiana*, *Sedum lineare*, *Hylotelephium erythrostictum*, *Iris germanica*, *Paeonia lactiflora*, *Hemerocallis dumortieri*, *Hosta plantaginea*, *Iris lacteal* (Figure 1). *Cosmos bipinnata* obtained the highest MP value with 92%, and *Dianthus barbatus* take second place with 73%, that this 2 species were significant higher than (*P*<0.05) the other species. *Iris lacteal* obtained the lowest MP mean value with 14%, was significantlower (*P*<0.05) than *Physostegia virginiana*, *Hylotelephium erythrostictum*, *Cosmos bipinnata* and *Dianthus barbatus*.

**Figure 1.**
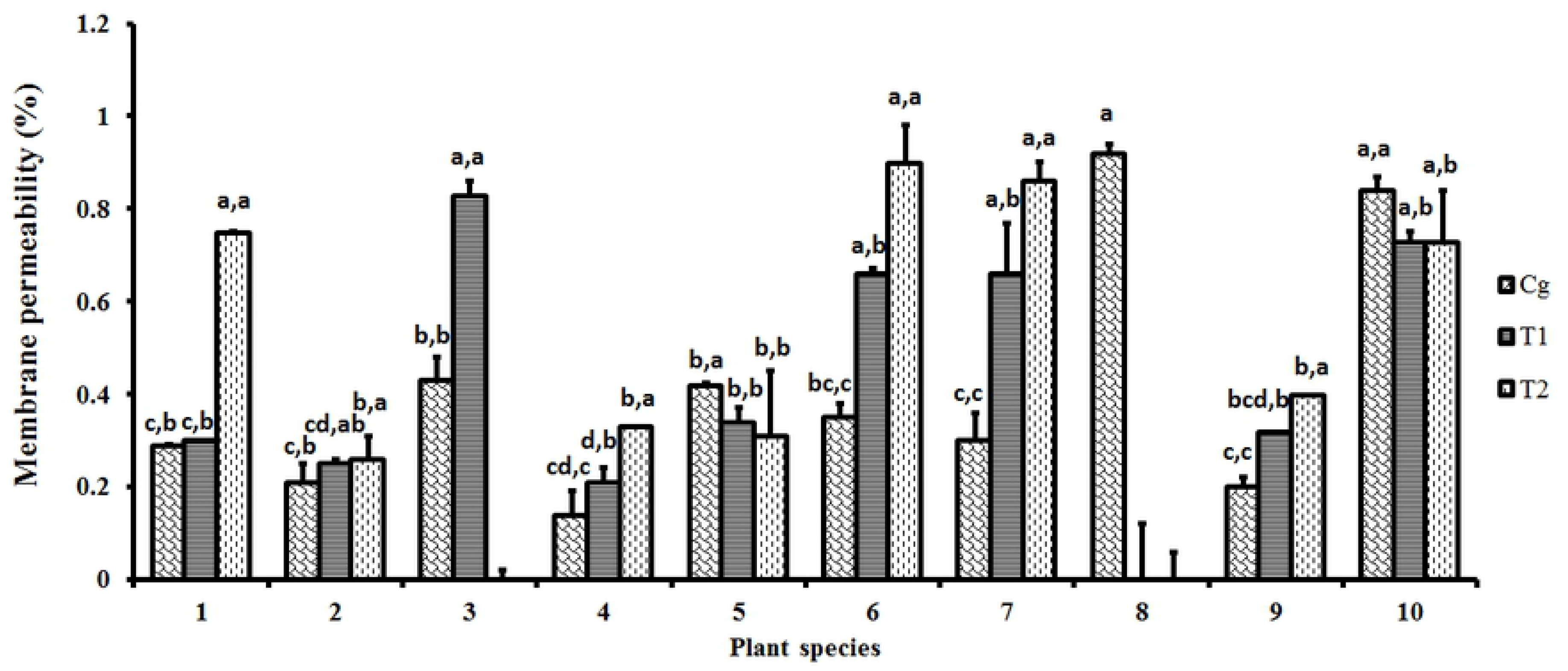
Membrane permeability changes of 10 species in the drought stress tests. Note: The histogram shows the mean values. *Cg* is short for control group (with normal watering condition); *T1* represents the “moderately drought stress”; *T2* represents the “strong drought stress”. The lowercase letter before the comma shows the statistical significance among different plant species, the lowercase letter behind the comma shows the statistical significance among *Cg*, *T1* and *T2*. Different lowercase letters indicate a significant difference at *P* < 0.05.

The cytomembrane is very sensitive to adverse condition and the plasmalemma would be stricken first by the drought stress (Yang *et al*., 2003). Plant species *Cosmos bipinnata* cannot live over the stress and dead in the moderate drought treatment (*T1*). When the seedlings meets *T1*, the MP mean value from high to low as *Physostegia virginiana*, *Dianthus barbatus*, *Sedum lineare*, *Iris germanica*, *Hosta plantaginea*, *Hylotelephium erythrostictum*, *Paeonia lactiflora*, *Hemerocallis dumortieri*, *Iris lactea*. The MP value of species *Physostegia virginiana* were the highest, was significantly different (*P*<0.05) with *Paeonia lactiflora*, *Hemerocallis dumortieri*, *Iris lactea*, *Hylotelephium erythrostictum*, *Hosta plantaginea*. The MP value of species *Physostegia virginiana*, *Iris lactea*, *Sedum lineare*, *Iris germanica* and *Hosta plantaginea* raised rapidly under *T1*. The MP of these 5 plants were significant higher (*P*<0.05) than those under *Cg*. This indicate these plant species’ cell may interrupted by the drought stress. On the contrary, *T1* reduced the MP values of species *Hylotelephium erythrostictum* and *Dianthus barbatus* significantly (*P*<0.05) compared with *Cg*. This at least suggests that the cells of these 2 plants were not further damaged. In addition, species *Paeonia lactiflora* and *Hemerocallis dumortieri* was still keep steady MP values.

Drought stress brought different influence on various plant species. Plant species *Physostegia virginiana* was dead during the strong drought treatment (*T2*). Most species got higher mean MP value under strong drought stress (*T2*) (Figure 1), except *Hylotelephium erythrostictum* and *Dianthus barbatus*. The MP mean values under *T2* from high to low as *Sedum lineare*, *Iris germanica*, *Paeonia lactiflora*, *Dianthus barbatus*, *Hylotelephium erythrostictum*, *Hosta plantaginea*, *Iris lactea*, *Hemerocallis dumortieri*. The MP values of species *Paeonia lactiflora*, *Iris lactea*, *Sedum lineare*, *Iris germanica* and *Hosta plantaginea* was significant higher (*P*<0.05) than those in the *T1*. The higher MP values, the more cytosol exosmosis, which suggests further damage to the cellular structure of these plants. Compared with other 4 species, *Hemerocallis dumortieri*, *Iris lactea*, *Hylotelephium erythrostictum*, *Hosta plantaginea* obtained lower MP mean values, and there were significant differences (*P*<0.05) with them.

### 4.3 Total chlorophylls contents

Usually, when the leaves of plants lost too much water, it would impede the chlorophyll production and may even resolve the chlorophyll (Yang *et al*., 2004; Zhang *et al*., 2003). That’s because the ROS (^1^O_2_, O_2_ ^−^ and ⋅OH) can make the lipid peroxidation directly and indirectly, thus damage the chlorophyll, and reduce the ratio of chlorophyll a and b (Zhang and Tan, 2001).

According to Figure 2, the Chl content of plant species from high to low as *Hemerocallis dumortieri*, *Paeonia lactiflora*, *Hosta plantaginea*, *Dianthus barbatus*, *Physostegia virginiana*, *Cosmos bipinnata*, *Iris germanica*, *Iris lactea*, *Hylotelephium erythrostictum* and *Sedum lineare*, under control treatment (*Cg*). The highest Chl content was belong to the species *Hemerocallis dumortieri* with 47.69 mg/g**⋅**_FW_, that was significant higher (*P*<0.05) than *Iris lactea*, *Hylotelephium erythrostictum*, *Sedum lineare*, *Iris germanica*. Species *Sedum lineare* obtained the lowest Chl content, with 6.06 mg/g**⋅**_FW_, that was significant lower (*P*<0.05) than the other plants species except *Hylotelephium erythrostictum*.

**Figure 2.**
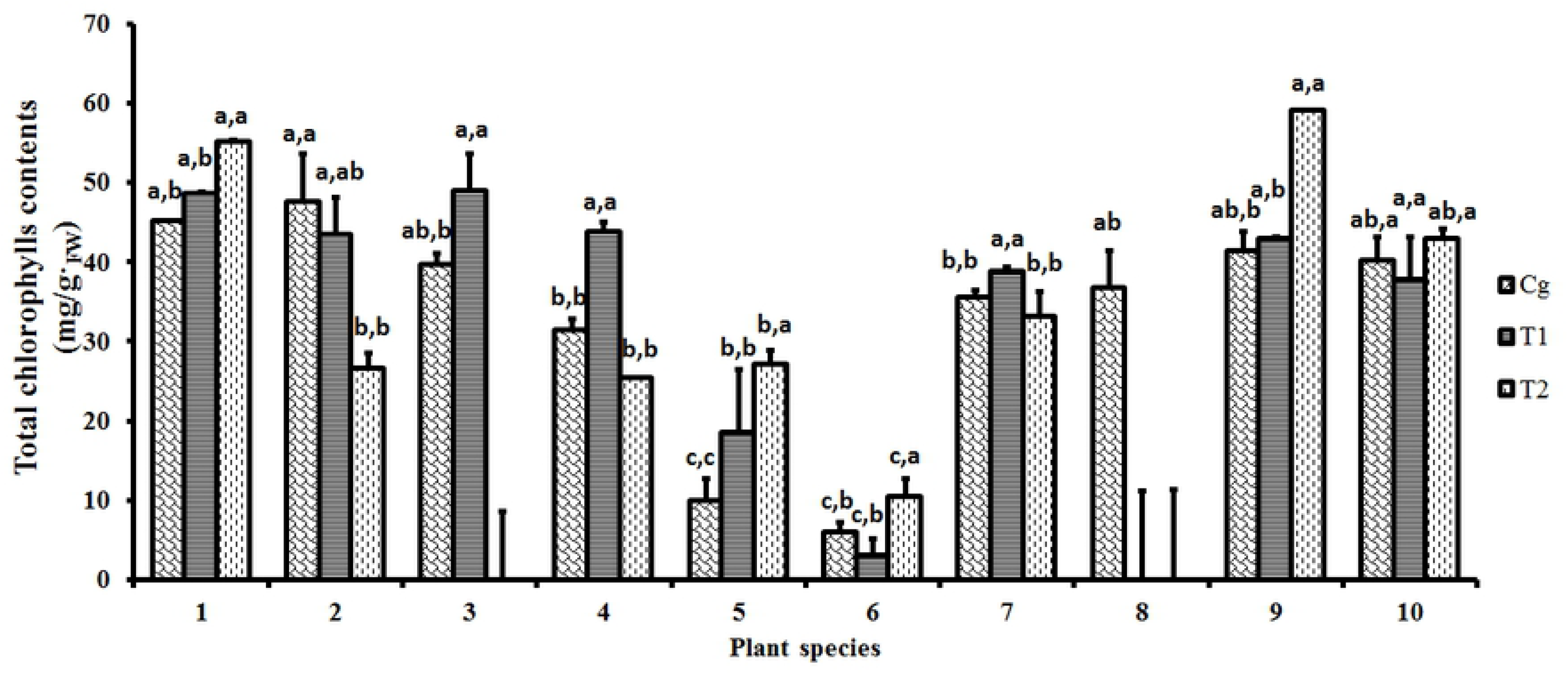
Total chlorophylls contents changes of 10 species in the drought stress tests. Note: The histogram shows the mean values. *Cg* is short for control group (with normal watering condition); *T1* represents the “moderately drought stress”; *T2* represents the “strong drought stress”. The lowercase letter before the comma shows the statistical significance among different plant species, the lowercase letter behind the comma shows the statistical significance among *Cg*, *T1* and *T2*. Different lowercase letters indicate a significant difference at *P* < 0.05.

Changes of Chl content in plants caused by drought stress. The Chl content of plant species from high to low as *Physostegia virginiana*, *Paeonia lactiflora*, *Iris lactea*, *Hemerocallis dumortieri*, *Hosta plantaginea*, *Iris germanica*, *Dianthus barbatus*, *Hylotelephium erythrostictum*, *Sedum lineare* under moderate drought treatment (*T1*). The Chl content of *Physostegia virginiana* was the highest with 49.07 mg/g**⋅**_FW_. The *Sedum lineare* obtained the lowest Chl content with 3.11 mg/g**⋅**_FW_, that was significant lower (*P*<0.05) than the other 8 surviving plants. There were 6 species’ Chl content increased include *Paeonia lactiflora*, *Physostegia virginiana*, *Iris lactea*, *Hylotelephium erythrostictum*, *Iris germanica* and *Hosta plantaginea* during *T1*. And *Hemerocallis dumortieri*, *Sedum lineare*, *Dianthus barbatus*’ Chl content was reduced by the *T1*.

The Chl content of plants from high to low as *Hosta plantaginea*, *Paeonia lactiflora*, *Dianthus barbatus*, *Iris germanica*, *Hylotelephium erythrostictum*, *Hemerocallis dumortieri*, *Iris lacteal* and *Sedum lineare*, under strong drought stress (*T2*). The Chl content of species *Hosta plantaginea* was the highest, with 59.11 mg/g**⋅**_FW_, that have significant difference (*P*<0.05) with species *Hemerocallis dumortieri*, *Iris lacteal*, *Hylotelephium erythrostictum*, *Sedum lineare* and *Iris germanica*. The lowest Chl content was belonging to *Sedum lineare*, that was significant lower than the other 7 surviving plants. The Chl content of species *Paeonia lactiflora*, *Hylotelephium erythrostictum*, *Sedum lineare* and *Hosta plantaginea* was increased during *T2* compared with *T1*. And such the 4 species were significant higher (*P*<0.05) than those under *T1* and *Cg*. That indicates the chloroplasts these 4 plant species were not harmed by drought stress. The species *Dianthus barbatus* maintained a steady trend of change of Chl content among *Cg*, *T1*and *T2*. From the perspective of Chl content, these plants mentioned above have good resistance to drought stress.

### 4.4 Superoxide dismutase activity

The superoxide dismutases (SODs) constitute a first line of defense at the cellular level (Noctor and Foyer, 1998) against ROS (Alscher *et al*., 2002). SODs are typically classified into three different groups depending on the prosthetic metal present in the active site, and they are designated as CuZn-SODs, Mn-SODs and Fe-SODs (Fridovich, 1986). McCord and Fridovich (1969) described principle of chemical reaction of SOD to eliminate the ROS as follows: O_2_ ^−^+O_2_ ^−^+2H^+^→O_2_ +H_2_O_2_. Since the SOD exists in peroxisomes, glyoxysomes, vacuoles, the nucleus, and the extracellular matrix, it plays a critical role to the drought tolerance (Molina-Rueda *et al*., 2013). The SOD activity reflects the plant species’ adaption to the environmental stress; the higher activity values, the stronger the adaption (Xu *et al*., 2014).

According to Figure 3, the SOD activity of plant species from high to low as *Cosmos bipinnata*, *Hemerocallis dumortieri*, *Iris germanica*, *Hosta plantaginea*, *Iris lactea*, *Physostegia virginiana*, *Dianthus barbatus*, *Hylotelephium erythrostictum*, *Paeonia lactiflora* and *Sedum lineare* under control treatment (*Cg*). Species *Cosmos bipinnata*’s SOD activity was the highest with 0.94 U**⋅**mg^−1^., that have significant difference (*P*<0.05) with species *Paeonia lactiflora*, *Hylotelephium erythrostictum* and *Sedum lineare*. Specific environment influence SOD activity, Chen *et al* (2009) researched the SOD of *Cosmos bipinnata*, a higher SOD activity obtained with high temperature (50 ℃, 15 min) extract by phosphate buffer (pH= 7.8, 2 hours). Meanwhile, species *Sedum lineare*’s SOD activity was the lowest only with 0.04 U**⋅**mg^−1^., that was significant lower (*P*<0.05) than the other 10 plant seedlings.

**Figure 3.**
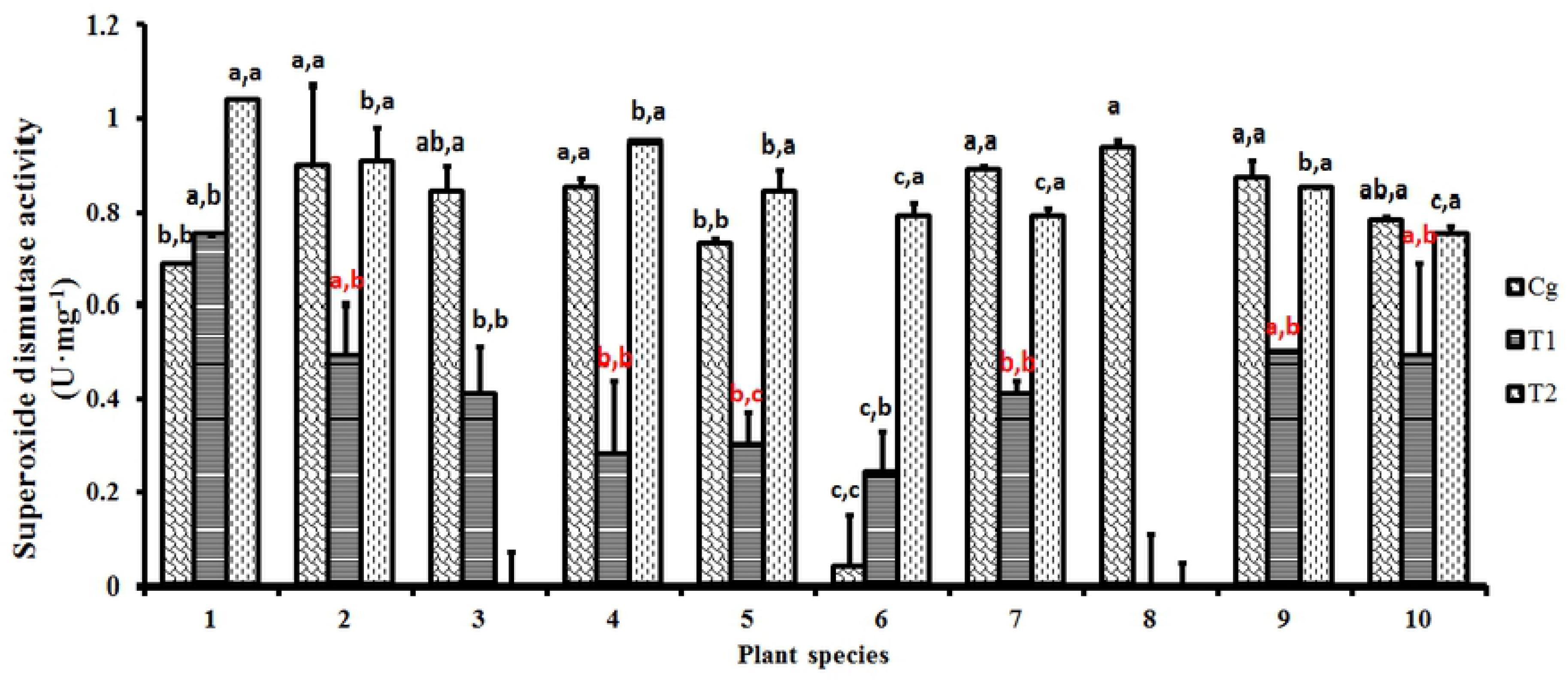
Superoxide dismutase activity changes of 10 species in the drought stress tests. Note: The histogram shows the mean values. *Cg* is short for control group (with normal watering condition); *T1* represents the “moderately drought stress”; *T2* represents the “strong drought stress”. The lowercase letter before the comma shows the statistical significance among different plant species, the lowercase letter behind the comma shows the statistical significance among *Cg*, *T1* and *T2*. Different lowercase letters indicate a significant difference at *P* < 0.05.

SOD activity is very sensitive to the drought environment (Ding *et al*., 2015; Wu *et al*., 2013; Zhang *et al*., 2014). The SOD activity of plant species from high to low as *Paeonia lactiflora*, *Hosta plantaginea*, *Hemerocallis dumortieri*, *Dianthus barbatus*, *Physostegia virginiana*, *Iris germanica*, *Hylotelephium erythrostictum*, *Iris lactea* and *Sedum lineare*, under the moderate drought treatment (*T1*). In there, the SOD activity of species *Hemerocallis dumortieri* and *Dianthus barbatus* was both 0.49 U**⋅**mg^−1^, and spices *Physostegia virginiana* and *Iris germanica* was both 0.41 U**⋅**mg^−1^. The species *Paeonia lactiflora* was significant higher (*P*<0.05) than *Physostegia virginiana*, *Iris lactea*, *Hylotelephium erythrostictum* and *Iris germanica*. Meanwhile, species *Sedum lineare*’s SOD activity was also the lowest only with 0.24 U**⋅**mg^−1^., that was significant lower (*P*<0.05) than the other 9 surviving plant seedlings. Species *Hemerocallis dumortieri*, *Iris lactea*, *Hylotelephium erythrostictum*, *Hosta plantaginea* and *Dianthus barbatus*’s SOD activity reduced rapidly, that have significant different (*P*<0.05) with those under *Cg*. This indicates such 5 species’ SOD activity interrupted by the drought. There are exceptions, the SOD activity of species *Paeonia lactiflora* and *Sedum lineare* was increased caused by drought.

The SOD activity of these plants that survived during strong drought stress (*T2*) was further enhanced compared with those under *T1*. The SOD activity of these plant seedlings from high to low as *Paeonia lactiflora*, *Iris lactea*, *Hemerocallis dumortieri*, *Hosta plantaginea*, *Hylotelephium erythrostictum*, *Sedum lineare*, *Iris germanica* and *Dianthus barbatus*. In there, species *Sedum lineare* and *Iris germanica* was both 0.79 U**⋅**mg^−1^. Species *Paeonia lactiflora*’s SOD activity was the highest with 1.04 U**⋅**mg^−1^, that was significant higher (*P*<0.05) than the other 8 surviving plant seedlings. Plants seedlings’ SOD activity was enhanced under strong drought, which indicates there is more protective enzyme produced by plants to eliminate ROS.

### 4.5 Peroxidase activity and Ascorbate peroxidase activity

Peroxidase is a kind of antioxidant enzymes that it could scavenge and decompose ROS (Mohammadi *et al*., 2016). The mechanism of POD scavenging ROS was described as follows: RH_2_+H_2_O_2_→2H_2_O+R, then H_2_O_2_ were transformed into H_2_O thoroughly (Asada, 1992; Qin *et al*., 2005). Survila *et al* (2016) demonstrates that increased peroxidase activity increases permeability of the leaf cuticle. When Chen *et al* (2017) and Wu *et al* (2016) researched the drought resistance of Cucumber and dendrobium moniliforme, they found that POD increased activity helps alleviate oxidative damage.

According to the Figure 4, POD activity kept a relatively high level in the *Cg*. *Hosta plantaginea* achieved 246.33 U**⋅**g^−1^**⋅**min^−1^, and it was significantly higher (*P*<0.01) than the other species except *Hemerocallis dumortieri*. The lowest species was *Physostegia virginiana*, only 14.36 U**⋅**g^−1^**⋅**min^−1^. When the seedlings suffered with *T1*, all the species’ POD activity reduced sharply. The highest POD activity was *Hemerocallis dumortieri* with 22.08 U**⋅**g^−1^**⋅**min^−1^, thus it have no significant difference with the other species. In addition, the higher peroxidase enzyme activity was obtained under drought stress and can be attributed to the plant defense mechanisms against free radical formation resulting from water defcit (Ruppenthal *et al*., 2016). In the case of *T1*, *Dianthus barbatus*, *Iris germanica* and *Iris lactea* showed stronger POD activity compared with the others. This indicates these 3 seedlings have better drought resistance to the moderately drought stress. Similarly, when the plants suffered with the strong drought stress, the POD activity of seedlings were further decreased. POD activity of *Paeonia lactiflora*, *Iris germanica* and *Dianthus barbatus* showed more sensitivity to the strong drought stress. They were significantly lower (*P*<0.01) than the *T1*.

**Figure 4.**
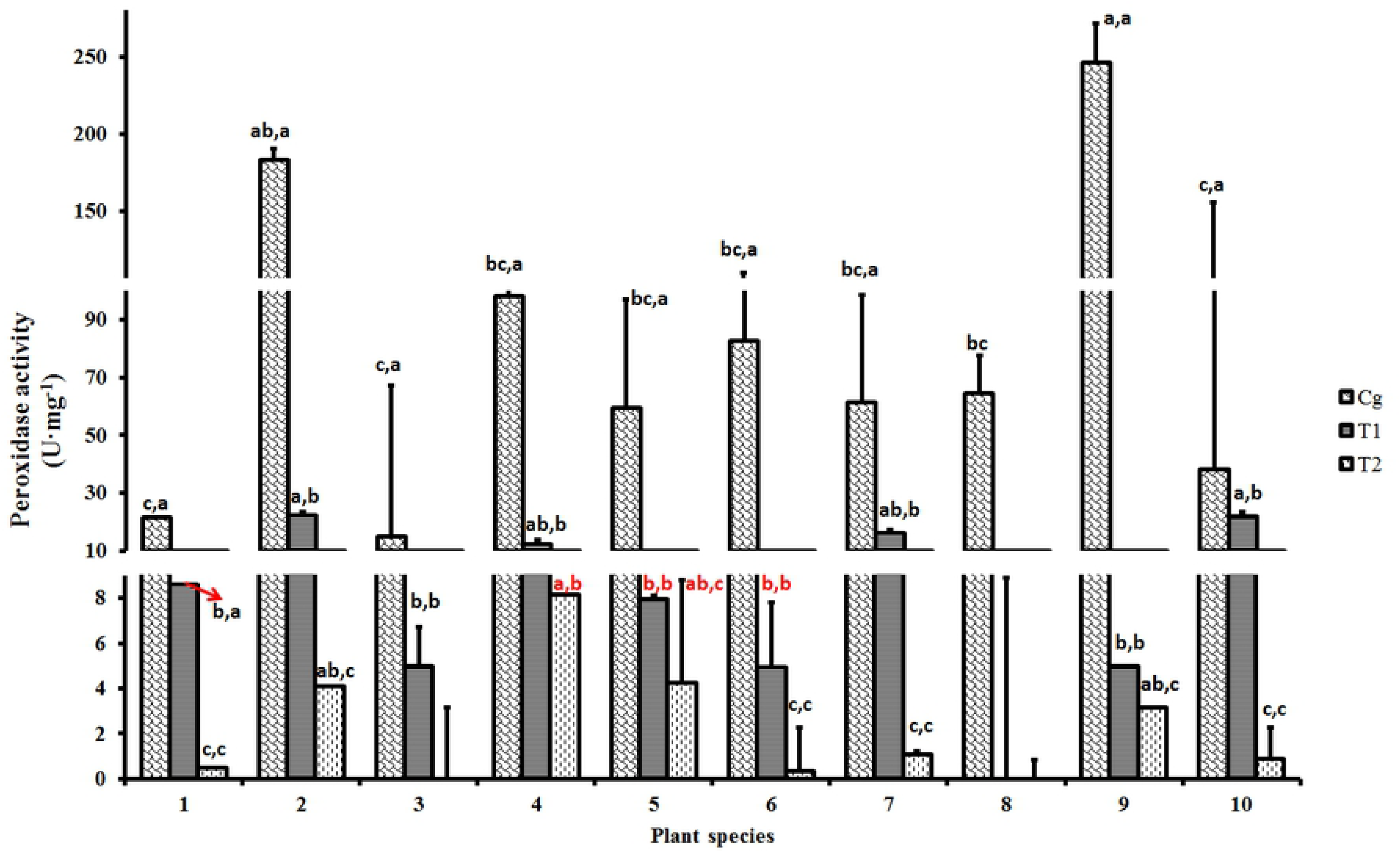
Peroxidase activity changes of 10 species in the drought stress tests. Note: The histogram shows the mean values. *Cg* is short for control group (with normal watering condition); *T1* represents the “moderately drought stress”; *T2* represents the “strong drought stress”. The lowercase letter before the comma shows the statistical significance among different plant species, the lowercase letter behind the comma shows the statistical significance among *Cg*, *T1* and *T2*. Different lowercase letters indicate a significant difference at *P* < 0.05.

In general, POD activity of all the species showed an obviously decreasing trend when the seedlings suffered with drought stress. The reason for the POD activity reduction in this research may be the different protective enzymes work as a whole eliminating the ROS. Such that when the SOD eliminates O_2_ ^─^, the H_2_ O_2_ would increase during the physiological reaction processes. However, with the drought stress enhancing, the reduction of POD activity impeded scavenge H_2_O_2_ (Sun *et al*., 2003). This indicates the plant seedlings defensive system of eliminating the ROS was too weak.

Ascorbate peroxidase belongs to the class I heme-peroxidases and is found in most eukaryotes including higher plants (Anjum *et al*., 2016). Ascorbate peroxidase exists as isoenzymes and plays an important role in the metabolism of H_2_O_2_ in higher plants (Asada, 1992; Shigeoka *et al*., 2002; Sukweenadhi *et al*., 2017). Meanwhile, AsAPOD also have the ability to scavenge the ROS (Li *et al*., 2013). That means the higher AsAPOD activity in plants resulted in faster removal of H_2_O_2_, which leads to alleviation of oxidative damage (Wu and Xia, 2006).

According to the Table 7, the highest values of AsAPOD activity was happened in the *Cg*, that they were significantly higher (*P*<0.05) than *T1* and *T2*. The AsAPOD activity is very similar to the change of POD activity. In the *Cg*, *Paeonia lactiflora* obtained the highest value as 17.86 U**⋅**min^−1^**⋅**g^−1^_FW_. The others present an equally same trend, that the values were during the 9.50 U**⋅**min^−1^**⋅**g^−1^_FW_ to 7.06 U**⋅**min^−1^**⋅**g^−1^_FW_. When the seedlings suffered with *T1*, the AsAPOD activity languished from the drought stress observably, and all of them were significantly lower (*P*<0.01) than the *Cg*. All the plant’s AsAPOD activity was less than 1.00 U**⋅**min^−1^**⋅**g^−1^_FW_, except for *Paeonia lactiflora* in the *T1*. All the plant’s AsAPOD activity shows a further decline in the *T2*, none of 0.50 U**⋅**min^−1^**⋅**g^−1^_FW_. The record was still *Paeonia lactiflora* in the *T2*. The change of AsAPOD activity was intense during the drought stress test, thus indicating AsAPOD was sensitive to the drought stress. As for AsAPOD activity of the 10 plant species, *Paeonia lactiflora* showed a better resistance to ROS.

**Figure 5.**
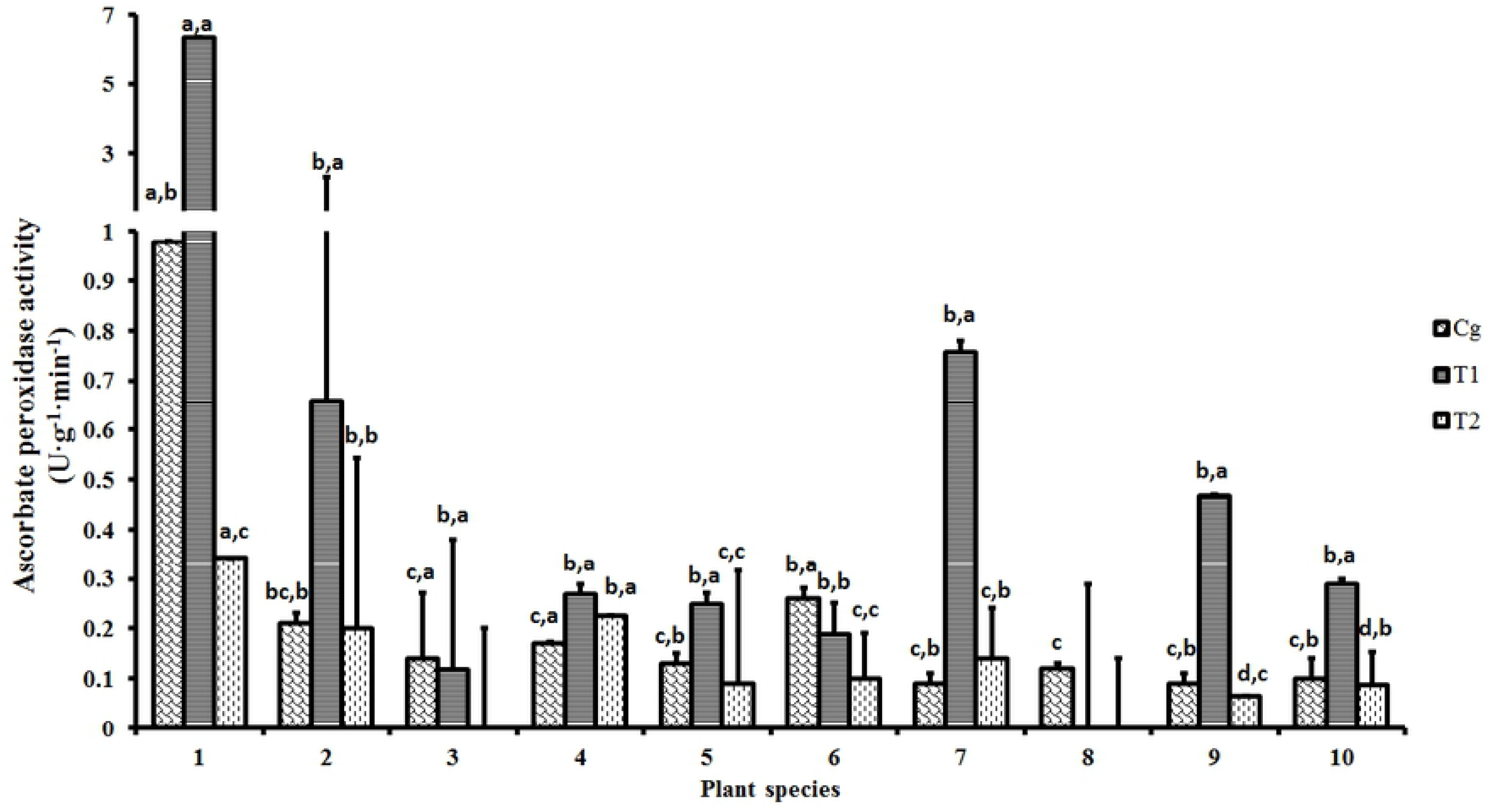
Ascorbate peroxidase activity changes of 10 species in the drought stress tests. Note: The histogram shows the mean values. *Cg* is short for control group (with normal watering condition); *T1* represents the “moderately drought stress”; *T2* represents the “strong drought stress”. The lowercase letter before the comma shows the statistical significance among different plant species, the lowercase letter behind the comma shows the statistical significance among *Cg*, *T1* and *T2*. Different lowercase letters indicate a significant difference at *P* < 0.05.

### 4.6 Assessment of drought resistance to the plants

The drought resistance of botany is a complex process, because the physiological and biochemical of plants changed as the ROS changed. A subordinate function method can be used for the assessment of drought resistance to the plants. The method’s essence was to apply fuzzy mathematics to the resistance indexes, such as DRP, Chl, SOD, POD and AsAPOD in this research. According to the formula of subordinate function, the results showed as following Table 4.

**Table 4.**
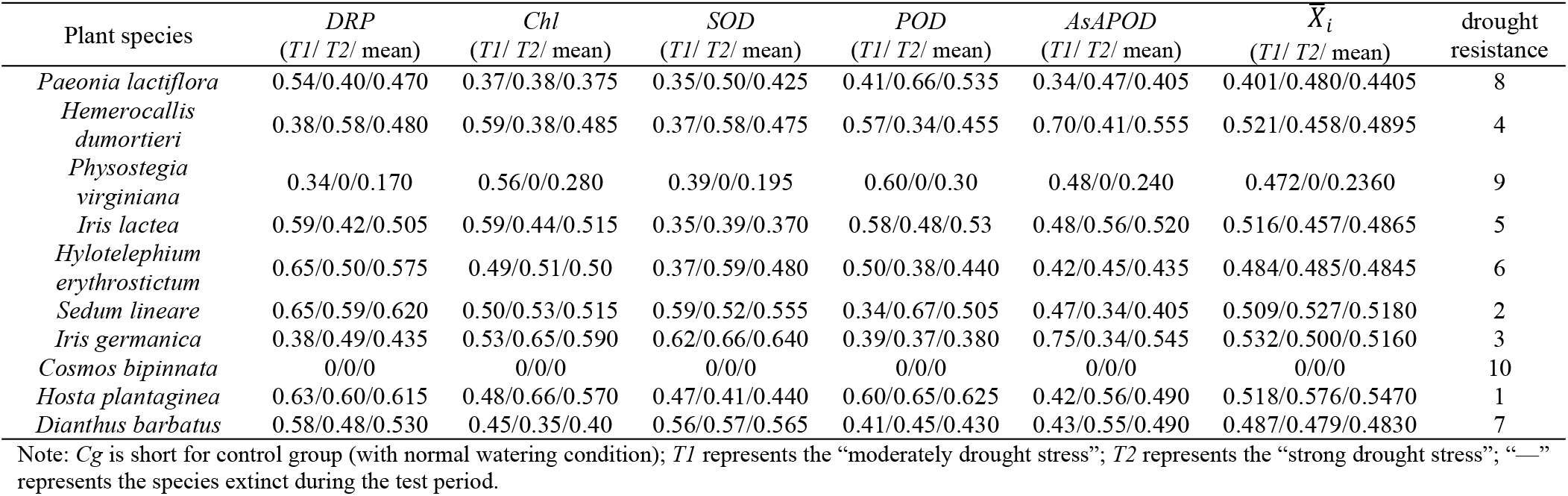
The drought resistance under the drought stress test

| Plant species | DRP<br>(T1/ T2/ mean) | Chl<br>(T1/ T2/ mean) | SOD<br>(T1/ T2/ mean) | POD<br>(T1/ T2/ mean) | AsAPOD<br>(T1/ T2/ mean) | $\bar{X}_i$<br>(T1/ T2/ mean) | drought<br>resistance |
| --- | --- | --- | --- | --- | --- | --- | --- |
| *Paeonia lactiflora* | 0.54/0.40/0.470 | 0.37/0.38/0.375 | 0.35/0.50/0.425 | 0.41/0.66/0.535 | 0.34/0.47/0.405 | 0.401/0.480/0.4405 | 8 |
| *Hemerocallis dumortieri* | 0.38/0.58/0.480 | 0.59/0.38/0.485 | 0.37/0.58/0.475 | 0.57/0.34/0.455 | 0.70/0.41/0.555 | 0.521/0.458/0.4895 | 4 |
| *Physostegia virginiana* | 0.34/0/0.170 | 0.56/0/0.280 | 0.39/0/0.195 | 0.60/0/0.30 | 0.48/0/0.240 | 0.472/0/0.2360 | 9 |
| *Iris lactea* | 0.59/0.42/0.505 | 0.59/0.44/0.515 | 0.35/0.39/0.370 | 0.58/0.48/0.53 | 0.48/0.56/0.520 | 0.516/0.457/0.4865 | 5 |
| *Hylotelephium erythrostictum* | 0.65/0.50/0.575 | 0.49/0.51/0.50 | 0.37/0.59/0.480 | 0.50/0.38/0.440 | 0.42/0.45/0.435 | 0.484/0.485/0.4845 | 6 |
| *Sedum lineare* | 0.65/0.59/0.620 | 0.50/0.53/0.515 | 0.59/0.52/0.555 | 0.34/0.67/0.505 | 0.47/0.34/0.405 | 0.509/0.527/0.5180 | 2 |
| *Iris germanica* | 0.38/0.49/0.435 | 0.53/0.65/0.590 | 0.62/0.66/0.640 | 0.39/0.37/0.380 | 0.75/0.34/0.545 | 0.532/0.500/0.5160 | 3 |
| *Cosmos bipinnata* | 0/0/0 | 0/0/0 | 0/0/0 | 0/0/0 | 0/0/0 | 0/0/0 | 10 |
| *Hosta plantaginea* | 0.63/0.60/0.615 | 0.48/0.66/0.570 | 0.47/0.41/0.440 | 0.60/0.65/0.625 | 0.42/0.56/0.490 | 0.518/0.576/0.5470 | 1 |
| *Dianthus barbatus* | 0.58/0.48/0.530 | 0.45/0.35/0.40 | 0.56/0.57/0.565 | 0.41/0.45/0.430 | 0.43/0.55/0.490 | 0.487/0.479/0.4830 | 7 |

Note: Cg is short for control group (with normal watering condition); T1 represents the “moderately drought stress”; T2 represents the “strong drought stress”; “—” represents the species extinct during the test period.

Through the calculation, the drought resistance of 10 plant species was very clear. *Iris germanica* was the best in drought resistance. *Cosmos bipinnata* and *Physostegia virginiana* died when they suffered with moderately drought stress and strong drought stress, respectively. *Paeonia lactiflora* survived the weakest drought resistance. The order of plants’ resistance to the drought resistance is as follow: *Hosta plantaginea > Sedum lineare > Iris germanica > Hemerocallis dumortieri > Iris lactea >Hylotelephium erythrostictum > Dianthus barbatus > Paeonia lactiflora > Physostegia virginiana > Cosmos bipinnata*.

## 5 Conclusions

The drought stress disturbed the plant growth, the MP, Chl, SOD, POD and AsAPOD have been changed compared to the control group. The result shows *Cosmos bipinnata* and *Physostegia virginiana* died after having suffered with moderately drought stress and strong drought stress, respectively. Although other species survived, the individual variation was huge especially for physiological and biochemical index. *Hemerocallis dumortieri*, *Iris lactea* and *Hosta plantaginea*’s MP had little change when they lived in the normal water condition and suffered with drought stress. The change of SOD activity is sensitive to drought stress. The most of the species (except *Paeonia lactiflora* and *Sedum lineare*) showed a lower SOD activity during moderately drought stress compared to the sufficient soil water condition and strong drought stress condition. The changes of plants’ POD activity and AsAPOD activity are very similar: when drought stress enhanced, the activity of protect enzyme reduced.

As a kind of local flowering plant, *Hosta plantaginea* shows good drought resistance. It is suggested to apply to the roof greening in Beijing and other northern cities in China widely. *Cosmos bipinnata* and *Physostegia virginiana* are too sensitive to drought stress, so it is not suggested to apply them to the roof greening, especially in the arid region. Other species can be applied to the roof greening for creating the landscape with no limitations. In order to increase the plants’ rate of survival and control the cost of roof greening, those plants with good drought resistance should be considered first.

## Acknowledgements

This work was supported by Young Core Plan of BJAST (Beijing Academy of Science and Technology) (No. 201528), Beijing Natural Science Foundation (No. 8142017) and National Natural Science Foundation of China (No. 41475133). Qin Fen, Business Development Manager for China office from Power Systems Research, a USA marketing research company in engine-driven industry, helped the manuscript preparation.

